# Elephant endotheliotropic herpesvirus is omnipresent in elephants in European zoos and an Asian elephant range country

**DOI:** 10.1101/2020.11.16.386011

**Authors:** Tabitha E. Hoornweg, Willem Schaftenaar, Gilles Maurer, Petra B. van den Doel, Fieke M. Molenaar, Alexandre Chamouard-Galante, Francis Vercammen, Victor P.M.G. Rutten, Cornelis A.M. de Haan

## Abstract

Elephant endotheliotropic herpesviruses (EEHVs) are a group of evolutionary divergent herpesviruses that may cause acute, often lethal, hemorrhagic disease (EEHV-HD) in young elephants. Although EEHV was first discovered over 20 years ago, its prevalence in different elephant populations is still largely unknown, partially due to the lack of readily available, sensitive serological assays. In order to improve diagnostic tools for the detection of EEHV infections and to obtain insight in its spread among elephants, we developed novel ELISAs focusing on EEHV1A gB and gH/gL as antigens. Performance of the ELISAs was assessed using sera taken from 41 European zoo elephants and 69 semi-captive elephants from Laos, one of the Asian elephant range countries. Sera from all (sub)adult animals tested (≥5 years of age) showed high reactivity with both gB and gH/gL, whereas reactivity towards the antigens was generally lower for sera of juvenile animals (1 > 5 years). Only one (juvenile) animal, which was sampled directly after succumbing to EEHV-HD, was found to be seronegative for EEHV. The two other EEHV-HD cases tested showed low antibody levels, suggesting that all three cases died upon a primary EEHV infection. Direct comparison with another EEHV-specific ELISA previously used in two large serosurveys, showed that EEHV prevalence was underestimated before, likely due to aberrant folding of the antigen used. In conclusion, our study suggests that essentially all (semi-)captive (sub)adult elephants in European zoos and in Laos carry EEHV, and that young elephants with low antibody levels are at risk of dying from EEHV-HD.

**Importance:** Over the last 30 years, nearly 20% of all Asian elephants born in Western zoos succumbed to acute hemorrhagic disease caused by elephant endotheliotropic herpesvirus (EEHV-HD). Yet, the prevalence of EEHV in captive and wild elephant populations is still largely unknown, mainly due to the lack of readily available, sensitive serological assays. For this study two highly sensitive EEHV-specific ELISAs were developed. Using these assays, it was shown that nearly all elephants tested were seropositive for EEHV, with highest antibody levels detected in (sub)adult elephants. In contrast, antibody levels in EEHV-HD cases were very low or non-detectable. Lack of antibodies may thus be a risk factor for developing severe disease. As the novel ELISAs are low-tech in nature, these assays may easily be disseminated to local laboratories in zoos and elephant range countries in order to determine EEHV serostatus of individual animals or complete herds and (wild) populations.

## Introduction

Elephant endotheliotropic herpesviruses (EEHVs) are responsible for a significant mortality rate observed in captive Asian elephants (*Elephas maximus*), and increasingly in African elephants (*Loxodonta africana*) in western zoos (1-3). Over the last decades, nearly 20% of all Asian elephant calves born in Western zoos succumbed to EEHV-HD (1). Furthermore, multiple reports showed that the disease affects both captive and wild elephants in the Asian elephant range countries (4-6). How widespread the virus is, and its prevalence in free-living populations, has yet to be established.

EEHVs are a group of evolutionary divergent herpesviruses, of which eight types have been identified so far (3). Four (sub)types (EEHV-1A, -1B, -4 and -5) naturally infect Asian elephants, while the other types (EEHV-2, -3, -6, and -7) infect African elephants. Like other members of the herpesvirus family, EEHVs cause latent infections in their host species. Healthy elephants can intermittently shed one or multiple EEHV subtypes as part of the natural infection cycle (7). It has been suggested that primary infections of young elephants, commonly aged between 1 to 8 years, may lead to acute, often lethal EEHV-Haemorrhagic Disease (EEHV-HD), which is particularly observed for EEHV-1A in Asian elephants (3, 8).

Currently, both the assessment of EEHV infections of individuals animals and the assessment of herd prevalence are largely performed using PCR on blood samples (5, 6, 9). Latent EEHV infections can only be detected upon reactivation, hence detection of EEHV prevalence by PCR requires longitudinal sampling over an extended period of time. Since PCR is relatively costly, and requires specialized equipment and trained personnel, the availability of this technique is limited in zoo laboratories and many elephant range countries (10). The virus-specific immune response is expected to remain present for the lifetime of the host (11). Therefore, serology would be the most convenient and cost effective method to assess EEHV prevalence in elephant populations.

To date, two different serological assays have been described. Van den Doel et al. (2015) designed an EEHV-specific ELISA with *E*.*coli*-expressed EEHV1A glycoprotein B (gB) as the antigen (12). This assay has been used to test two large cohorts of elephant sera and showed that 37% of elephants from Western zoos (12) and 42% of captive elephants in Thailand (13) were seropositive for EEHV. Although it was shown that EEHV infections were relatively wide-spread, 24% of the elephants with PCR-confirmed EEHV infection were designated EEHV seronegative in this assay (12), indicating that this assay probably underestimates EEHV seropositivity. Recently, a second serological assay was described by Fuery et al. (8), who used four different EEHV proteins genetically fused to luciferase and expressed in mammalian cells as antigenic fractions. It was shown that 82% of all elephants tested (N=24, from four different North American zoos), including 100% of the adult animals (≥ 15 years of age; N=10), were EEHV seropositive (8).

The herpesvirus gB, glycoprotein H (gH) and glycoprotein L (gL) are conserved between all herpesviruses and essential for host cell entry. The fusion protein gB naturally forms a homotrimer (14). A heterodimer of gH and gL acts as an activator of gB and may function in receptor binding (15, 16). Both gB and the gH/gL dimer are important targets of the herpesvirus-specific humoral immune response (17-20), hence interesting for use in EEHV-specific serological assays. Although expression of recombinant EEHV gB has been reported (8, 12), purification has only been achieved from bacterial cultures. Expression of recombinant EEHV gH/gL has not been described thus far.

The current study aimed to improve diagnostic tools for EEHV infection in elephants, thereby focussing on gB and gH/gL as antigens, and, by use of these tools, to obtain novel insight into the spread of EEHV among elephants. We describe the successful purification of secreted recombinant EEHV1A gB and gH/gL produced in mammalian cells. Both gB and gH/gL were strongly recognized by antibodies present in elephant sera, indicating their value as antigens in EEHV-specific ELISAs. We subsequently showed that all subadult (between 5 and 15 years of age) and adult elephants (≥ 15 years of age), living under human care in either European zoos (N=34) or an Asian elephant range country (Laos, N=69), were seropositive for EEHV. Furthermore, all three EEHV-HD cases tested had low or non-detectable EEHV-specific antibody titres, indicating that these animals experienced primary infections as previously suggested (8).

## Results

### Production and purification of recombinant EEHV1A gB, gH/gL and gL

The DNA constructs for the production of soluble recombinant EEHV1A glycoproteins gB, gH and gL were designed as described in the Materials and Methods section. The resulting glycoproteins are schematically depicted in Fig. 1A. For all EEHV glycoproteins, the native signal sequences were exchanged for the Gluc signal sequence, transmembrane and cytoplasmic regions (gB and gH) were removed, and a StrepTag (ST) or HisTag (HT) was added at the protein C-terminus. Upon transfection of the gB-ST, gH-ST, gL-ST constructs and the IAV HA-ST control construct into HEK293T cells, production of proteins of the expected size was observed (Fig. 1B cell lysate fractions and Table 1). Of these proteins, gB-ST, gL-ST and HA-ST were secreted into the supernatant (Fig. 1B secreted fractions), while no secretion of gH-ST could be observed. When EEHV1A gH and gL were co-expressed, intracellular production and secretion of both proteins was observed as expected (Fig. 1B) (21-24).

**Table 1.** Predicted molecular weights of the protein constructs used in the study.

| **Construct** | **Predicted MW (kDa)<sup>a</sup>** | **Predicted N-linked glycosylation sites<sup>b</sup>** | **Predicted MW, N-linked glycosylation included (kDa)<sup>c</sup>** |
| --- | --- | --- | --- |
| gB-ST | 80,4 | 6 | 95,4 |
| gH-ST | 84,6 | 8 | 104,6 |
| gL-ST | 34,8 | 4 | 44,8 |
| gL-HT | 29,9 | 4 | 39,9 |
| gH-ST/gL-ST | 119,4 | 8/4 | 149,4 |
| gH-ST/gL-HT | 114,5 | 8/4 | 144,5 |
| IAV HA-ST | 42,5 | 4 | 52,5 |

<sup>a</sup>Molecular weights (MWs) were predicted using the online expasy PI/Mw tool
(https://web.expasy.org/compute_pi/).
<sup>b</sup>The number of N-linked glycosylation sites were predicted using the NetNGlyc server
(http://www.cbs.dtu.dk/services/NetNGlyc/).
<sup>c</sup>To estimate the MW of the glycosylated proteins, a MW of 2.5 kDa was added for each predicted N-linked glycan.

**Figure 1.**
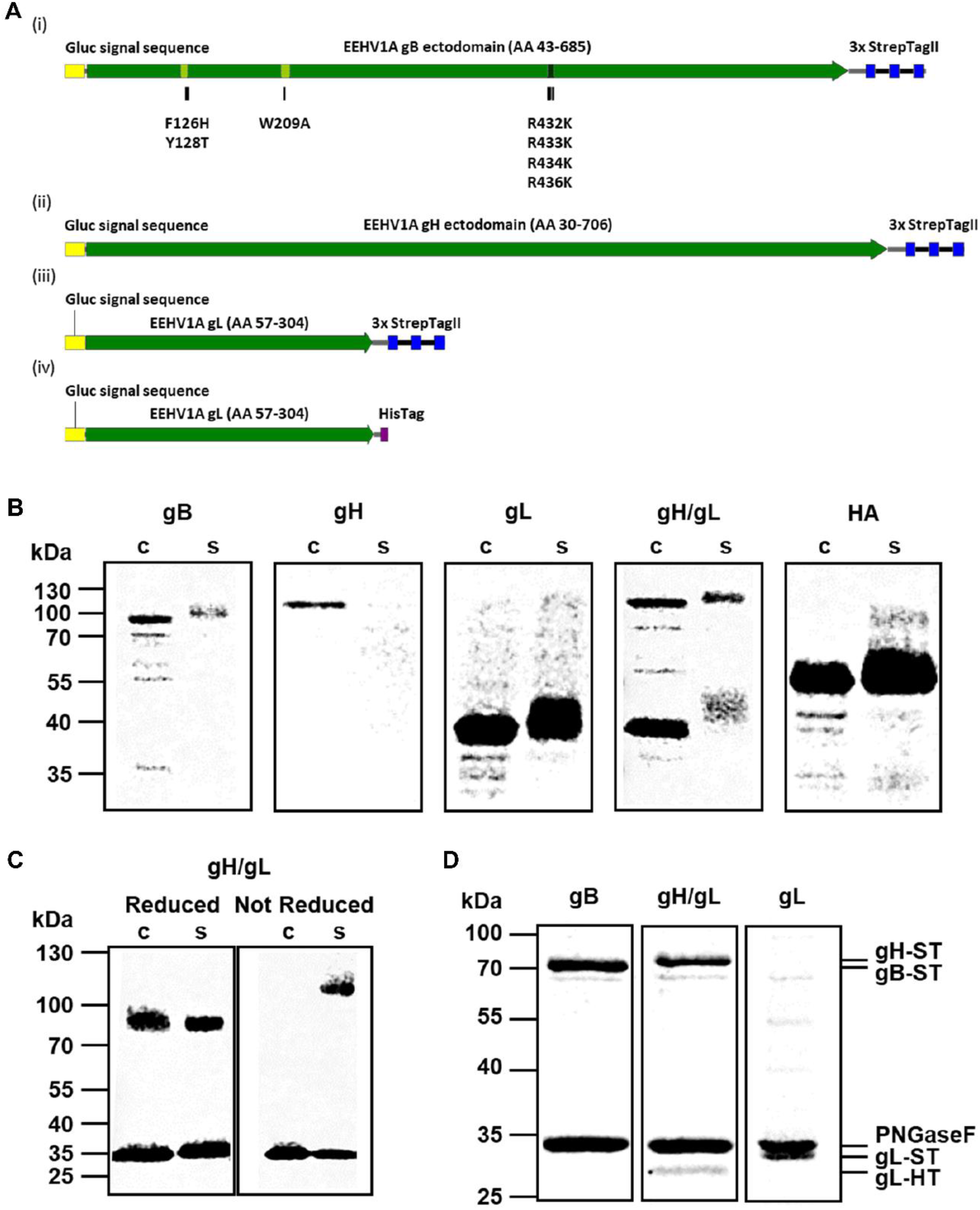
Production of recombinant EEHV1A gB, gH/gL and gL using the HEK293T expression system. (A) Schematic representation of the recombinant (i) gB-ST, (ii) gH-ST, (iii) gL-ST and (iv) gL-HT proteins. Amino acid mutations introduced into the two putative fusion loops (light green) and furin cleavage site (dark green) of gB are indicated as black bars. (B) Representative Western Blots showing the cell lysate (c) and secreted (s) fractions of recombinant EEHV1A gB, gH, gL, gH/gL and the IAV HA control produced in HEK293T cells. (C) Western Blot of the cell lysate (c) and secreted (s) fractions harvested from a gH-ST/gL-ST co-transfection subjected to SDS-PAGE under reducing and non-reducing conditions. Proteins were deglycosylated by PNGaseF prior to electrophoresis. (D) Gelcode Blue-stained purified EEHV1A gB-ST, gH-ST/gL-HT and gL-ST. Samples were deglycosylated by PNGaseF prior to electrophoresis to facilitate analysis. Molecular mass markers are indicated on the left side of the gels.

Previous studies on HCMV revealed that gH and gL are covalently linked by disulphide bonds to form a dimer (25). Amino acid alignments of the HCMV and EEHV gH or gL sequences indicated that the cysteines involved in the formation of these disulphide bonds are conserved between HCMV and EEHV (Supplementary Figure 1). To explore whether the EEHV1A gH/gL heterodimer is also linked by disulphide bonds, the cell lysate and secreted EEHV1A gH/gL fractions were subjected to SDS-PAGE under reducing and non-reducing conditions. Under reducing conditions, two individual protein bands representing gH (∼85 kDa) and gL (∼35 kDa) were visible in both fractions (Fig. 1C; reducing gel). Under non-reducing conditions, monomeric gL remained visible in both fractions, while monomeric gH was not visible. Instead, a protein band of approximately 120 kDa was observed in the secreted fraction, corresponding to the predicted molecular weight of the disulphide-linked gH/gL heterodimer (∼ 119 kDa) (Fig. 1C; non-reducing gel).

After confirmation of secretion of gB, gL and the gH/gL dimer into the supernatant, secreted proteins were subjected to affinity purification by use of their StrepTag. To prevent copurification of the gL monomer with the gH/gL heterodimer, the StrepTag in the gL construct was exchanged for a HisTag (Fig. 1A (iv)). Fig. 1D shows Gelcode Blue stained gels of the affinity purified gB-ST, gH-ST/gL-HT and gL-ST fractions. Samples were treated with PNGaseF to remove N-glycans in order to facilitate protein quantification. Clear protein bands of the expected molecular weights were visible in all three fractions with minor contamination. Copurification of gL-HT with gH-ST further supported the formation of stable gH/gL heterodimers (Fig. 1D). Intensity of Gelcode Blue staining showed that both proteins were present at equimolar concentrations, indicating that gH and gL interact in a 1:1 ratio.

### Development of EEHV-specific ELISAs using purified EEHV1A gB, gH/gL and gL proteins

The recombinant EEHV1A proteins were used for development of ELISAs to detect EEHV-specific antibodies in elephant sera. To define the optimal antigen concentrations for measurement of antibody titres, we used five individual elephant sera that had shown to differ to some extent in their reactivity with the EEHV proteins in preliminary experiments and a no serum control using dilution ranges of the different proteins produced. Four out of five elephant sera showed a clear dose-dependent response to the gB antigen (Fig. 2A), while the serum of one elephant and the no serum control were virtually unreactive. The same four elephant sera showed a dose-dependent response to gH/gL (Fig. 2B), albeit lower than for gB. Interestingly, none of the elephant sera were reactive towards gL (Fig. 2C).

**Figure 2.**
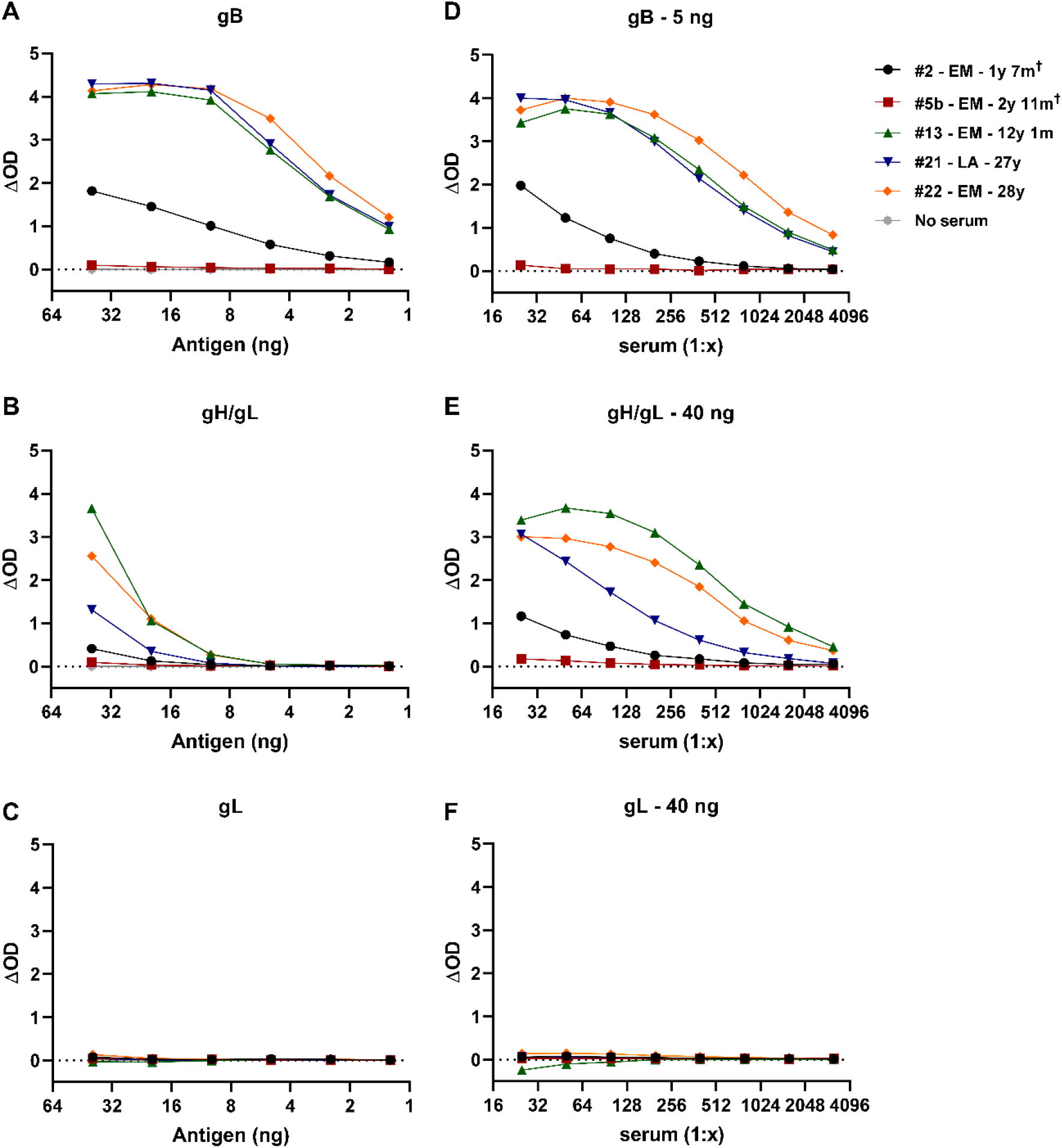
Antigen and serum dilution curves for the novel gB, gH/gL and gL ELISAs. Dilution ranges (40 – 1.25 ng/well) of (A) gB, (B) gH/gL and (C) gL. (D-F) Serum dilution ranges assayed using optimal antigen dilution determined in A-C, namely (D) 5ng gB, (E) 40 ng gH/gL and (F) 40 ng gL. Dilution ranges were performed either single or in duplo, and results were depicted as the ΔOD (signal in antigen-coated well – signal in uncoated well).

For gB and gH/gL, optimal protein concentrations for the detection of EEHV-specific antibodies were 5 respectively 40 ng/well, both in the linear range of the antigen dilution curve. As no reactivity towards gL was observed under the present assay conditions and coating of higher concentrations was not feasible because of limited protein quantities available, the highest concentration tested (40 ng/well) was used in further optimisation of the ELISA. To define the optimal serum concentrations to be used in the ELISAs, dilution ranges of the same five sera were tested in combination with the optimal antigen coating concentrations defined. Clear dilution-dependent responses were observed for gB (Fig. 2D) and gH/gL (Fig. 2E) for the same four sera that showed a dose-dependent response in the antigen dilution ranges, while the serum of one animal remained virtually unreactive at all dilutions tested. Again, none of the sera showed a response towards gL (Fig. 2F). A serum dilution of 1:100 was considered optimal for future ELISA testing as at this dilution high reactivity was combined with a large difference in reactivity between the different sera.

### Close to all tested zoo elephants are EEHV seropositive

Sera of 41 elephants from European zoos were tested using the novel EEHV ELISAs. The sample set included 28 single sera, seven longitudinal and one paired serum samples from Asian elephants, and five single sera from African elephants. Ages at time of sampling ranged from 1 year and 4 months to 56 years for the Asian elephants and from 27 to 36 years for the African elephants. Fig. 3A shows the results of the gB ELISA, sorted on elephant age. All elephants but one were found to have gB-specific antibodies, and all subadult and adult animals displayed high ΔOD levels (range: 2.0 - 4.2). Sera of six (of seven) juvenile Asian elephants showed a clear response towards gB, yet for most juveniles the ΔOD values were substantially lower than values obtained for the (sub)adult animals (range for positive animals: 0.6 - 4.2). Notably, sera of the three juvenile elephants sampled close to the time of death due to EEHV-HD showed low (#2 and #6) or non-detectable (#5) gB-specific antibody levels.

**Figure 3.**
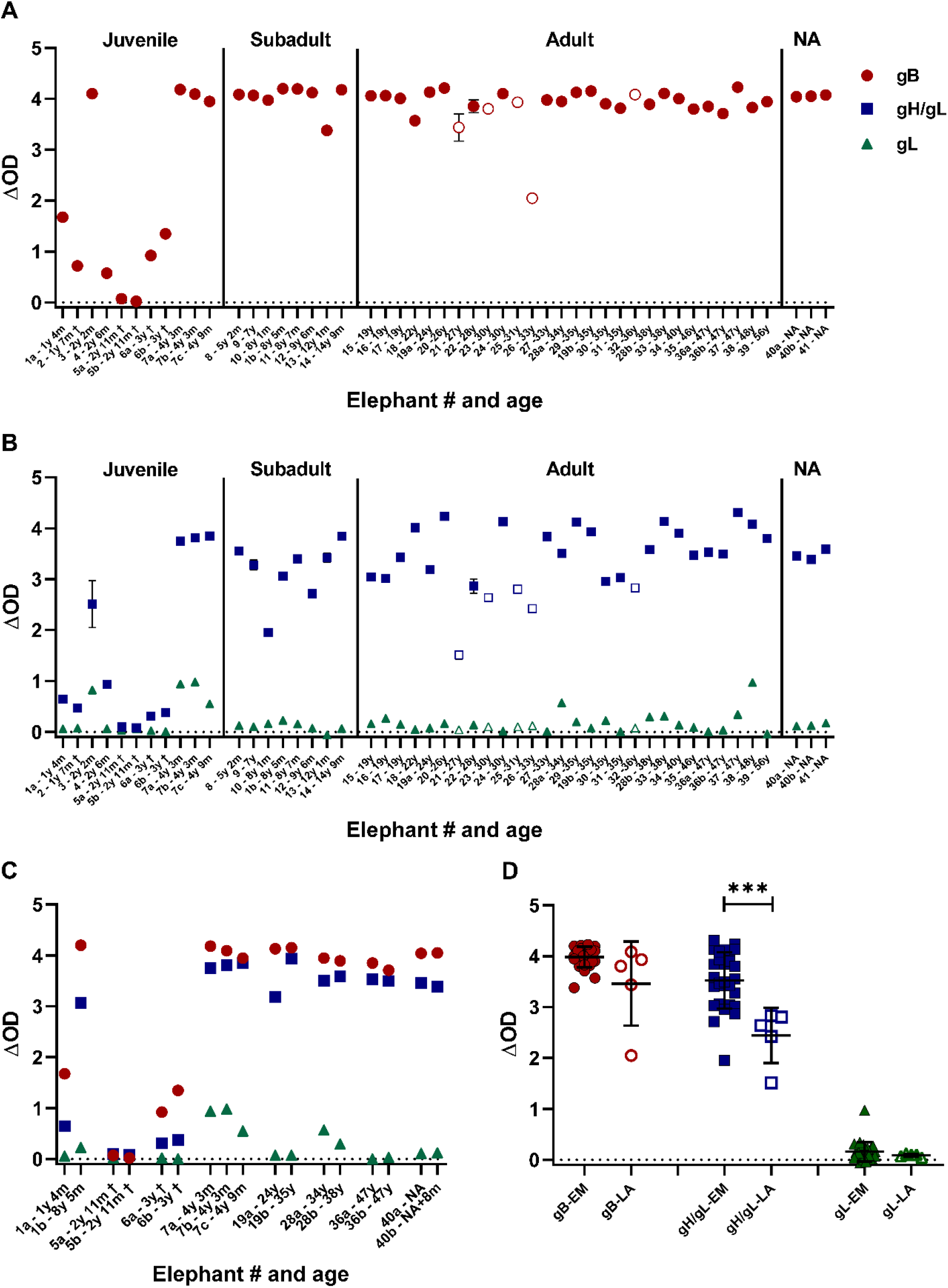
Assessment of EEHV seropositivity in elephants living in European zoos. ΔOD values of 50 elephant sera from 41 individual elephants obtained in the gB (A), gH/gL (B – blue squares) and gL (B – green triangles) ELISAs. Sera are identified by elephant number and age and ordered based on age at time of sampling on the x-axis. Longitudinal or paired samples of individual elephants are identified by the individual elephant number followed by a character (either a, b or c). Samples from Asian elephants (*Elephas maximus* = EM) are shown by closed symbols and those from African elephants (*Loxodonta Africana* = LA) by open symbols. The majority of samples were tested once, while 8 (gB), 10 (gH/gL) and 4 (gL) sera were assayed multiple times (range 2-10 times) with comparable results. For these samples, standard deviations are indicated, which are only visible when they exceed the size of the symbol used. (C) ΔOD values of the longitudinal or paired serum samples of eight individual elephants. (D) ΔOD values of individual (sub)adult elephants divided by elephant species. For elephants with longitudinal samples available, the serum of the last sampling date was selected for analysis. Significance was tested by Mann-Whitney-test: *** = p ≤ 0.001.

When the same serum set was tested in the gH/gL ELISA, comparable results were obtained (Fig. 3B, blue squares). All elephants, except the one juvenile elephant, were seropositive for the antigen, and ΔOD values were generally higher in (sub)adult animals (range: 1.5 - 4.3) than in the juvenile animals (range for positive animals: 0.3- 3.8). Again, gH/gL-specific antibody levels in the juveniles that succumbed to EEHV-HD were low (#2 and #6) or non-detectable (#5). Finally, sera from all animals were tested in the gL ELISA (Fig. 2B, green triangles). In line with the results during ELISA optimization, most animals were unreactive towards gL. Nevertheless, a low but specific response was noted for some animals.

The ΔOD values of the eight elephants for which longitudinal or paired samples were available (all included in Fig. 3A as well) are shown in Fig. 3C once more. Notably, a clear increase in ΔOD values against both gB and gH/gL was observed for animal #1, which was 1 year and 4 months old at first sampling and over eight years old at second sampling. Also, for animal #19, titres against gH/gL clearly increased during the 11 year sample interval, while titres against gB remained at similar (high) levels. For the other animals, ΔOD values remained relatively stable between samples, most likely as a result of the high reactivity already observed at first sampling (animal #7, 28, 36 and 40) and short term sampling intervals (animal #6). For animal #5 two different sample types (heparin and EDTA plasma), both collected directly post mortem, showed comparable results.

The above results show that the novel gB and gH/gL ELISAs identify the same animals as EEHV seropositive. Nevertheless, it was noted that the ΔOD values obtained in the gH/gL ELISA (Fig. 3B) were much more variable between animals than the values obtained in the gB ELISA (Fig. 3A). Comparison of the ΔOD values obtained for Asian and African elephants revealed a significant difference in the gH/gL, but not in the gB ELISA (Fig. 3D).

### Comparison of the novel gB ELISA to the previously published EEHV gB ELISA

The performance of the novel EEHV-specific gB ELISA was compared with that of a previously published EEHV-specific gB ELISA (12). Differences between both assays include the method through which the gB antigen is produced: *in casu* gB produced in mammalian cells or in *E*.*coli* respectively.

A randomly selected subset of sera from the European zoo cohort (20 sera from 17 Asian elephants and one serum from an African elephant) were assayed by us in both ELISAs. The ΔOD values are shown in Fig. 4A. Sera that gave rise to the highest ΔOD levels in the previously published ‘bacterial’ gB ELISA, showed relatively low ΔOD levels in the novel ‘mammalian’ gB ELISA. Vice versa, for many sera that tested highly positive in the ‘mammalian’ gB ELISA, ΔOD values around 0.0 were obtained in the ‘bacterial’ gB ELISA. Results obtained by both ELISAs were found to be negatively correlated with a R^2^-value of 0.42 (Fig. 4B).

**Figure 4.**
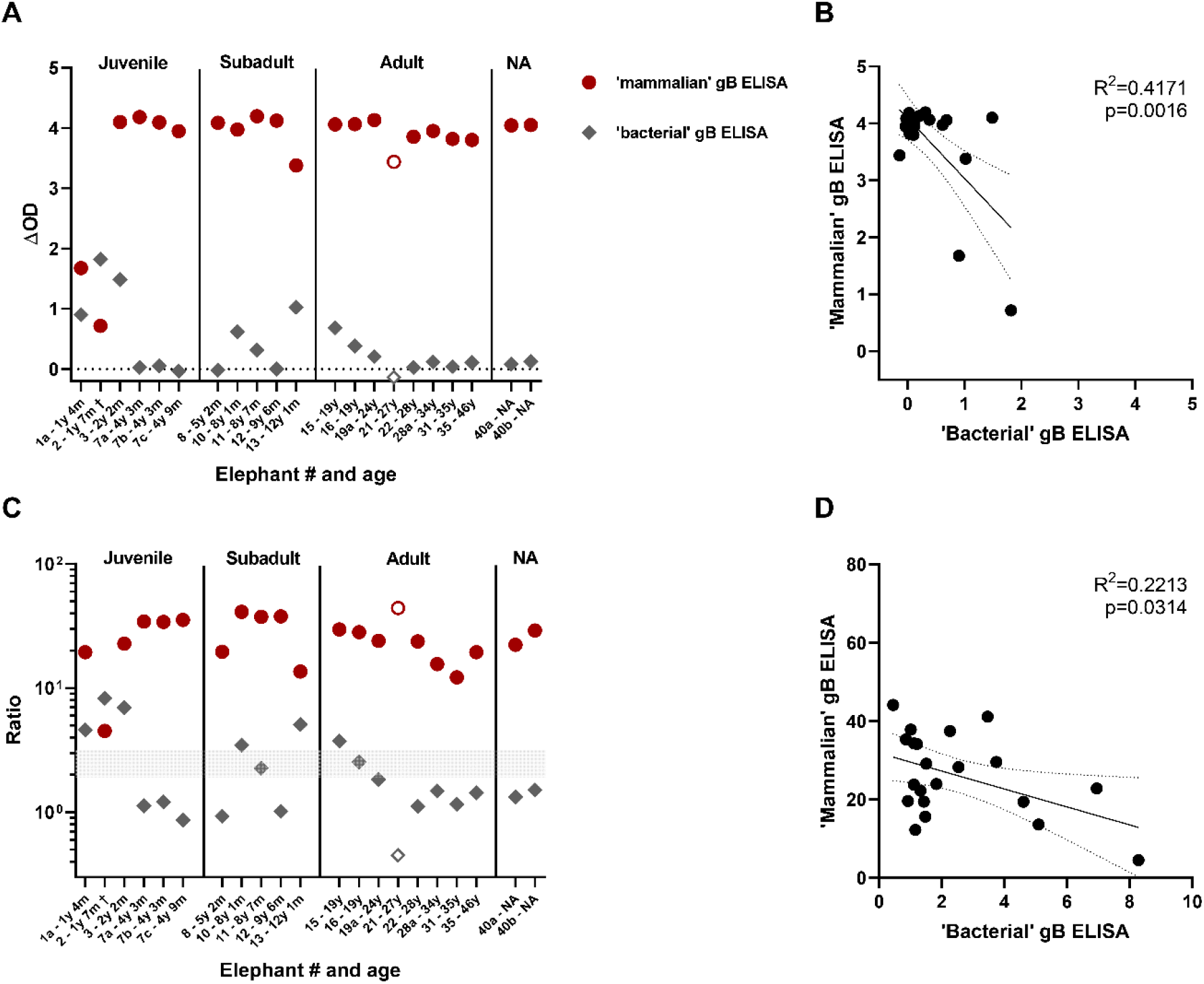
Comparison of the novel ‘mammalian’ gB ELISA to the ‘bacterial’ gB ELISA. (A) ΔOD values of 21 sera (of 18 individual elephants) tested using both the novel ‘mammalian’ gB ELISA (red circles) and a previously published ‘bacterial’ gB ELISA (grey diamonds) (12). Closed symbols indicate Asian elephants (EM, N=17) while open symbols African elephants (LA, N=1). Sera were tested at both 1:100 and 1:200 dilutions in the ‘bacterial’ gB ELISA; the most optimal (highest) ΔOD ratio is shown. (B) Correlation between the ΔOD values of the ‘mammalian’ and ‘bacterial’ gB ELISA shown in A. (C) Antigen-specific OD/background OD-ratio calculated for ELISAs. Sera were tested at both 1:100 and 1:200 dilutions in the bacterial gB ELISA; the most optimal (highest) ΔOD ratio is shown. Symbols as described in A. (D) Correlation between the antigen-specific OD/background OD-ratio’s shown in C.

Since seropositivity in the ‘bacterial’ gB ELISA is determined based on the signal/background-ratio instead of the ΔOD values (12), signal/background-ratios were calculated for both ELISAs and plotted in Fig. 4C. In line with previous publications, ratios above 3 were considered positive, ratios between 2 and 3 were considered borderline and ratios below 2 were determined to be negative (12, 13). By use of this cut-off all samples tested in the ‘mammalian’ gB ELISA were found seropositive for EEHV, while 6/21 (29%) and 2/21 (10%) of the samples were found to be either positive or borderline in the ‘bacterial’ gB ELISA, respectively. Also for these calculated ratios a negative correlation was observed between both ELISAs, albeit less significant and with a lower R^2^ value (Fig. 4D).

### Antisera generated against linear gB epitopes suggest altered epitope availability in gB produced in bacterial versus mammalian cells

A likely explanation for the observed differences between the ‘bacterial’ and ‘mammalian’ gB ELISAs would be different epitope availability between both recombinant gB proteins. To explore this hypothesis, rabbit sera generated against five different gB-specific peptides (Fig. 5A) were assessed for their abilities to recognize the mammalian and bacterial gB antigens in ELISA. In agreement with our hypothesis, differential recognition was indeed observed for the peptide-specific rabbit sera (Fig. 5B).

**Figure 5.**
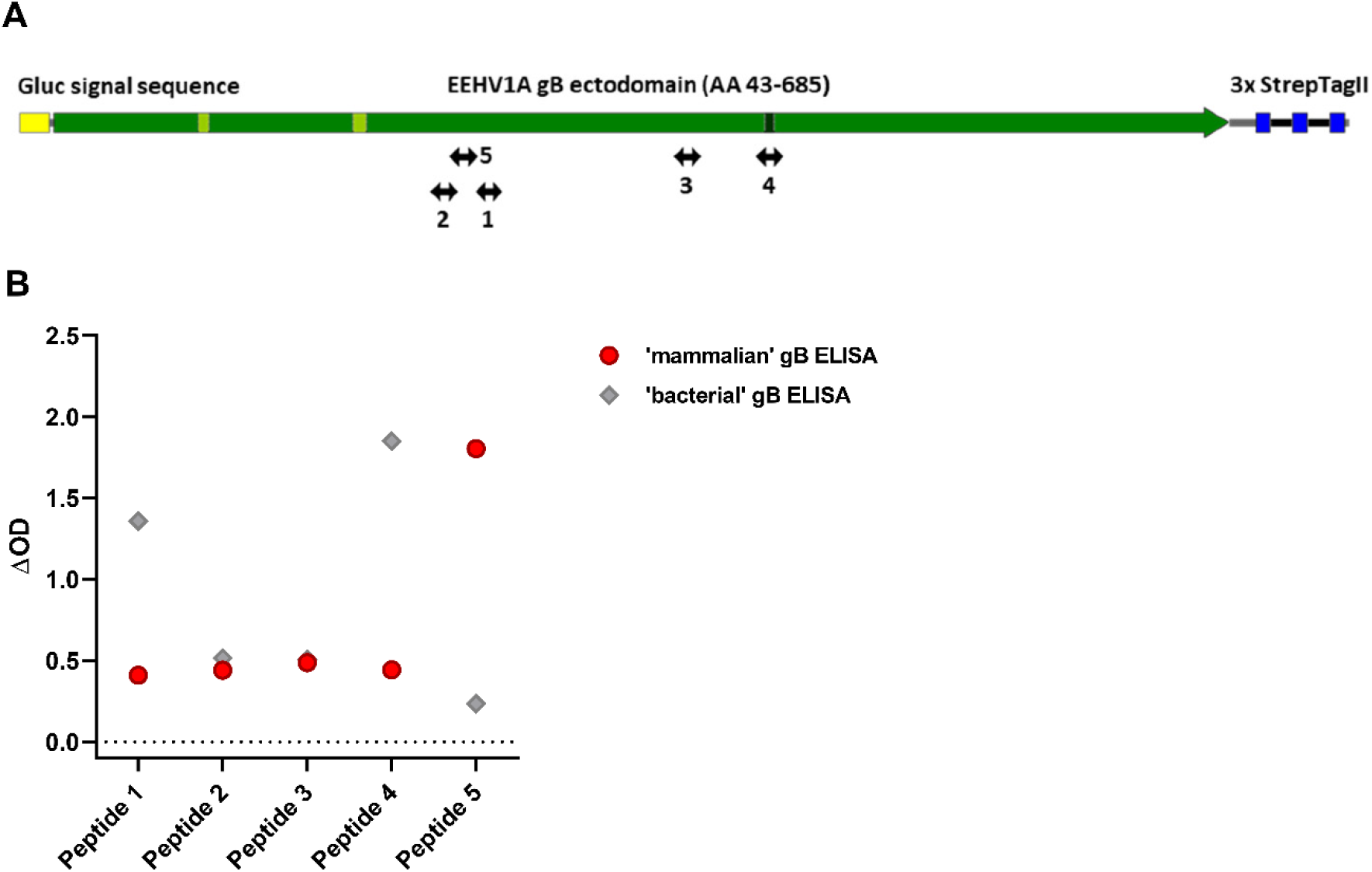
Reactivity of five EEHV1A gB peptide-specific rabbit antisera towards mammalian and bacterial produced gB. (A) Positions of the five gB peptides to which rabbit antisera were generated are indicated (12). (B) ΔOD values for the gB peptide-specific rabbit sera measured in the mammalian gB (red circles) and bacterial gB (grey diamonds) ELISAs. Sera were tested at least once in duplicate at a 1:100 dilution. Representative results are shown.

Mammalian-produced gB was best recognized by the serum generated against peptide 5, whereas bacterial-produced gB was primarily recognized by sera generated against peptides 1 and 4. For peptide 4 it must be noted that its epitope, which spans the furin cleavage site, was partially mutated in the ‘mammalian’ gB construct.

### All tested Laotian elephants under human care are seropositive for EEHV

After having shown that nearly all elephants in European zoos tested EEHV seropositive, we aimed to explore the EEHV status of Asian elephants living in range countries. Sera from 69 elephants could be included for analysis, and all were found to be seropositive for EEHV, with ΔOD values ranging between 3.2 - 4.3 and 2.2 - 4.2 for gB and gH/gL, respectively (Fig. 6). We conclude that all the Laotian elephants tested are infected with one or multiple EEHV types.

**Figure 6:**
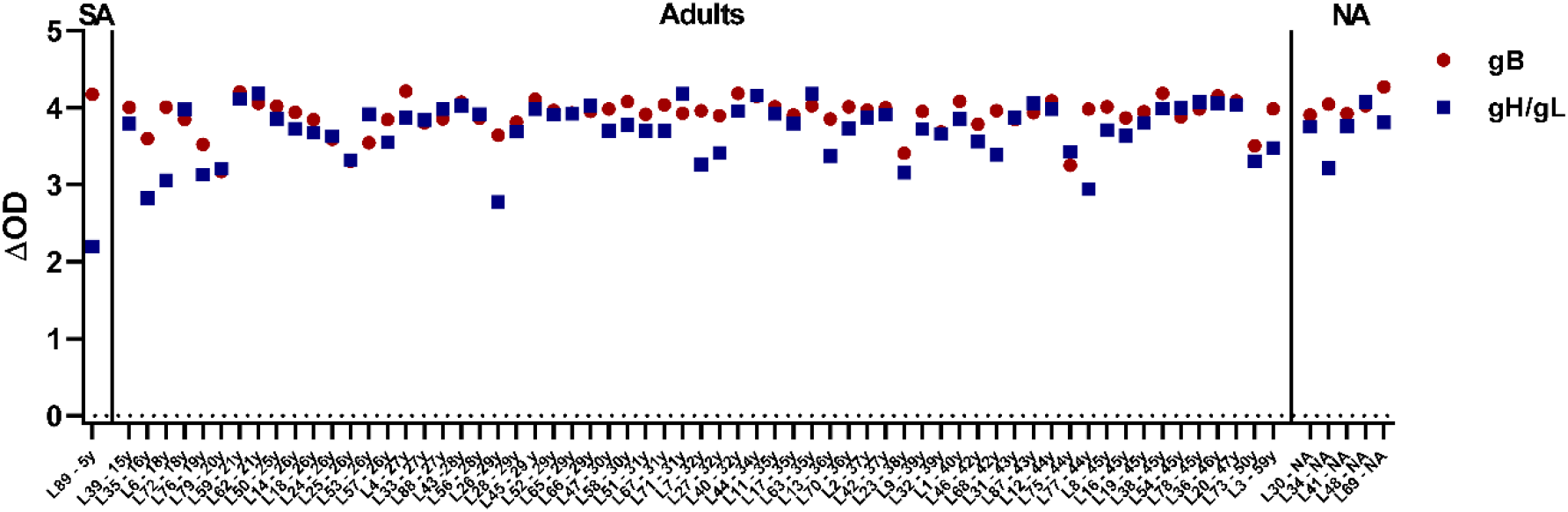
Results obtained in the gB and gH/gL ELISAs for 69 serum samples from individual Asian elephants living under human care in Laos. Samples are plotted based on age at sampling. SA= subadult, NA= information on age not available.

## Discussion

This study aimed at improving diagnostic tools for EEHV infection in elephants and obtaining novel insights into the spread of EEHV among elephants. Since the virus cannot be propagated *in vitro*, one of the major problems is availability of EEHV-specific antigens. Here, we report the successful production and purification of recombinant EEHV1A gB and gH/gL, and the subsequent development of EEHV glycoprotein-specific ELISAs. The novel ELISAs showed that 40 out of 41 screened elephants from European zoos were EEHV seropositive, and all 69 screened (semi-)captive elephants from Laos, one of the Asian elephant range countries.

The prevalence of EEHV positive animals in both cohorts included in this study was considerably higher than that previously reported in two other serosurveys (12, 13). In 2015, Van den Doel and coworkers showed that 37% of the European zoo elephants tested (N=125) were seropositive to EEHV (12), while in 2019 Angkawanish and coworkers reported that 42% of 994 Thai elephants living under human care were EEHV seropositive (13). Both serosurveys used the ‘bacterial’ gB ELISA, of which this study showed that it underestimates EEHV seropositivity as compared to the novel ELISAs (Fig. 4). This difference may be best explained by incorrect folding of *E. coli*-expressed gB due to a lack of mammalian-type glycosylation (27). Differential reactivity of antisera raised against a set of linear gB peptides is in agreement herewith (Fig. 5). The fact that the same animals were identified as EEHV seropositive by use of two different EEHV antigens, gB and gH/gL, while most animals were unreactive to a third EEHV antigen (gL), produced in the same manner and carrying the same protein tag as gB and gH/gL, further supported the validity of our results and excluded the possibility of non-specific binding to either the tag or possible contaminants introduced during protein production.

All (sub)adult Asian (N=98) and African (N=5) elephants tested in this study, including a large cohort from Laos (N=69), were seropositive to EEHV. High seroprevalence has been reported for many herpesvirus infections (28), however for EEHV similar seroprevalence (82%) has thus far only been reported for a relatively small number of elephants (N=24), living in four different North American zoos, of which all adult elephants (N=10) were EEHV seropositive (8). In line herewith, two studies that monitored EEHV shedding in a limited number of animals in two zoo herds for several months, showed that 72% (N=7 (29)) to 100% (N=6 (7)) of the elephants tested were infected with EEHV. Our study indicates that EEHV is widespread among Asian, and possibly also African elephants, both in zoos and in a range country (Laos) with potentially all adult animals infected with one or more EEHV subtypes. More studies are needed to establish whether this also holds true for free-ranging elephants.

In line with antibody responses to many human herpesvirus infections (17-20), both gB and gH/gL were identified as important targets of the EEHV-specific immune response, while gL, in absence of gH, was found not to be a major target of the natural EEHV-specific immune response. Interestingly, the anti-gH/gL response was found to be more variable between elephants than responses towards gB, which may be explained by elephants being infected with different EEHV types, and gH and gL being less conserved between the different EEHV species than gB (Table 2). In agreement herewith, anti-gB responses of Asian elephants, for which infections with EEHV subtype 1A are common, and of African elephants, which naturally carry other EEHV subtypes (30), were comparable in our ELISAs, while the responses towards gH/gL were significantly higher in Asian than in African elephants (Fig. 3D). Our data suggest that while the gB ELISA should be regarded as a pan-EEHV ELISA, gH/gL might be an interesting target for the development of subtype-specific EEHV ELISAs. The latter may be possible by comparing (end point) titers towards a set of subtype-specific gH/gL proteins, in analogy to comparative virus neutralization tests used for serotyping (human) flavivirus and orthohantavirus infections (31, 32).

**Table 2.**
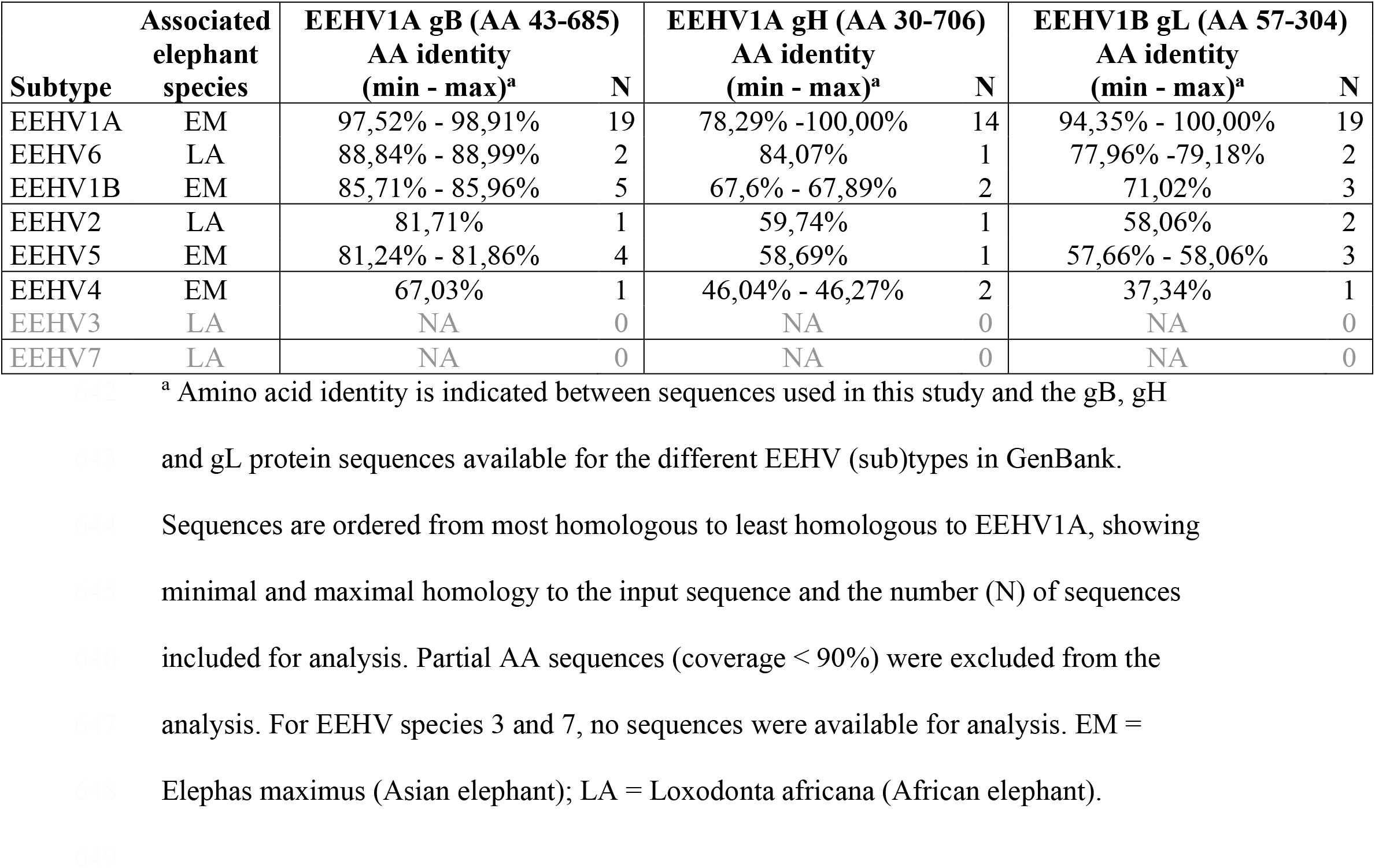
Amino acid identity between gB, gH and gL

Recently, Fuery and co-workers suggested that EEHV-HD was associated with a primary EEHV infection rather than virus reactivation (8). Within our European zoo cohort, sera of three juvenile animals that succumbed to EEHV1A-HD, collected during the EEHV-HD episode or directly post-mortem, were tested. EEHV-specific antibody levels in all three animals were low (N=2) or undetectable (N=1), which would be consistent with a primary EEHV infection rather than reactivation. This supports the hypothesis of EEHV-HD being the result of an uncontrolled primary infection.

Albeit based on a limited number of cases, the current study also suggests that not only completely seronegative animals, but also calves with low (maternal) antibody levels are at risk of developing EEHV-HD. Future studies are needed to discern how the levels, avidity and specificity of antibodies targeting gB and gH/gL are correlated with protection against severe disease, which is crucial knowledge for elephant management and future development of an effective EEHV vaccine. Pending these studies, these novel ELISAs may already be used for the detection of those young elephants that run the most risk of developing EEHV-HD. Furthermore, the relative low-tech nature of these novel ELISAs allows easy dissemination to local laboratories in Asian and African elephant range countries to determine the EEHV seroprevalence in larger cohorts of captive and, where blood access is possible, free-living elephants.

## Materials and Methods

### Expression of recombinant EEHV proteins

Codon-optimized cDNAs (Genscript) encoding EEHV1A gB (accession number AAN03667; residues 43-685), gH (accession number AGG16086; residues 30-706) and gL (accession number AGG16117; residues 57-304) were cloned into a pFRT expression plasmid (Thermo Fisher Scientific) in frame with sequences encoding the Gaussia luciferase (Gluc) signal sequence and a C-terminal StrepTag (ST) or HisTag (HT) as depicted in Fig. 1A. To increase expression levels, seven amino acid substitutions were introduced into the fusion loops (F126H, Y128T and W209A) and the furin cleavage site (R432K, R433K, R434K, R436K) of the gB construct.

HEK293T cells (ATCC CRL-3216) were cultured in DMEM (Gibco) supplemented with 10% Fetal Calf Serum (FCS; Bodinco), Penicillin (100 U/ml) and Streptomycin (100 μg/ml) (both from Lonza) at 37°C under 5% CO2. Individual plasmids or a combination of the gH and gL expression plasmids (4:1 ratio) were transfected into HEK293T cells using polyethylenimine (PolyScience) as described previously (33). A pFRT expression plasmid encoding the Influenza A virus (IAV) Haemagglutinin (H1) ectodomain coupled to a C-terminal Strep-Tag was used as a control (34). Five days post transfection cell culture media were harvested. Additionally the cells were lysed using Radioimmunoprecipitation assay (RIPA) buffer (150 mM NaCl, 1% Triton X-100, 0.5% sodium deoxycholate, 0.1% SDS, 50 mM Tris, pH 8.0) for 30 min at 4°C. Subsequently, cell culture media and lysates were centrifuged, supernatants were collected and subjected to sodium dodecyl sulfate-polyacrylamide gel electrophoresis (SDS-PAGE). When indicated, proteins were deglycosylated by PNGaseF (NEB) prior to gel electrophoresis. Proteins were transferred to a PVDF membrane using the Transblot Turbo System (BioRad), stained using a horseradish peroxidase (HRP)-conjugated monoclonal anti-StrepTag antibody (Iba) and detected by use of enhanced chemiluminescence (ECL) Western Blotting substrate (Pierce) and the Odyssey imaging system (LI-COR).

For purification, cell culture media containing the secreted EEHV glycoproteins were cleared of debris by low speed centrifugation and proteins were purified using Strep-Tactin Sepharose beads (Iba). Purified proteins were subjected to quantitative densitometry of GelCode Blue (Thermo Fisher Scientific)-stained protein gels containing bovine serum albumin (BSA) standards. Signals were imaged and analyzed with an Odyssey imaging system (LI-COR).

### Sera

Fifty serum samples of 41 individual elephants from 15 European zoos were used in this study. For most elephants (39/41), species, age at sampling and sex were known. All blood samples were taken aseptically from the ear or leg vein by zoo veterinary staff and sera were sent to our institutes for diagnostic purposes. Additionally, 77 sera from (semi-)captive elephants from Laos, previously described by Lassausaie et al. (26), were tested. The vast majority of these elephants were released into the forest for three to six months a year, where they could roam freely and interact with their wild counterparts. All samples were drawn in 2012 and for the vast majority of samples (72/77), sex and age at sampling were known. For both serum cohorts, elephants were classified in age groups according to Arivazhagan and Sukumar (35); Elephants <1 year old are considered as babies, those between 1 and 5 years are juveniles, and those between 5 and 15 years respectively ≥15 years as subadults and adults. Sera with high background values (OD-values >1 in the wells lacking antigen) were excluded from analysis (N=8, all from Laotian elephant cohort). The rabbit sera specific for gB peptides 1-5 used in this study were previously described by Van der Doel et al. (12). All sera were shipped cooled and stored at -20°C upon receipt.

### ELISAs

Purified recombinant EEHV1A gB, gH/gL or gL (diluted in PBS; 100 µl/well) were coated overnight on Nunc MaxiSorp high protein-binding capacity ELISA plates (ThermoFisher). Subsequently, plates were washed three times with PBST (PBS + 0.05% Tween-20) and incubated with blocking buffer (PBS containing 0.1% Tween20 and 3% BSA [w/v] for 2 hours. Next, plates were washed (three times with PBST) and 100 µl serum diluted in blocking buffer was added to the wells for 1 hour. Plates were washed (three times with PBST), and incubated with 100 µl HRP-conjugated recombinant Protein A/G (0.5 µg/ml diluted in blocking buffer; Pierce), previously reported to bind elephant IgG (36, 37), for 1 hour. After washing (three times with PBST), 100 µl/well TMB Substrate (Surmodics) was added, plates were incubated for 10 minutes in the dark, and the reaction was stopped by adding 100 µl 12,5% H2SO4. Optical density (OD) was measured at 450 nm in an EL-808 ELISA reader (BioTEK). For each sample, the antigen-specific signal (signal in wells containing antigen) and serum-specific background signal (signal in wells lacking antigen) were assessed simultaneously, and the ΔOD value (OD value antigen coated well – OD value uncoated well) was reported. Sera were excluded from analysis if OD-values in the wells lacking antigen were >1 (N=8, all from Laotian elephant cohort). The optimal antigen concentration was tested by antigen dilution (range 40 – 1,25 ng/well) using five elephant sera and a no serum control. After establishment of the optimal antigen concentration, the optimal serum dilution was determined using serum dilution ranges (1:25 – 1:3200) of the same elephant sera. To compare performance of the novel assay with the existing EEHV-specific ELISA using gB expressed in *E*.*coli* as antigen (12), a subset of 21 randomly selected sera were tested in both assays. The ‘bacterial’ gB ELISA was performed as previously described (12). Upon use of the rabbit anti-gB peptide sera to assess epitope accessibility in the novel ‘mammalian’ and the ‘bacterial’ gB ELISAs, HRPO-conjugated swine anti-rabbit IgG (1:1000, DAKO) was used as the conjugate.

## Acknowledgements

We would like to thank all zoos involved in this study for collection and sharing of elephant serum samples. In addition, we would like to thank dr. Jérôme Lassausaie and the staff of the Lao livestock department for sample collection from the Laotian elephants. Finally, we would like to thank the Royal Zoological Society of Antwerp, DierenPark Amersfoort Wildlife Fund, Abri voor Dieren Foundation, Animales Foundation, Utrecht University Fund, P. Zwart Foundation, A.A.M. Bijleveld Foundation, Marjo Hoedemaker Foundation, Named Fund Friends of VetMed, and many individual donors for their ongoing support of this research. Funders had no role in the study design, data collection and analysis, decision to publish, or preparation of the manuscript.

